# Meditation-induced effects on whole-brain structural and effective connectivity

**DOI:** 10.1101/2021.06.10.447903

**Authors:** Eleonora De Filippi, Anira Escrichs, Estela Càmara, César Garrido, Marti Sánchez-Fibla, Matthieu Gilson, Gustavo Deco

## Abstract

In the past decades, there has been a growing scientific interest in characterizing neural correlates of meditation training. Nonetheless, the mechanisms underlying meditation remain elusive. In the present work, we investigated meditation-related changes in structural and functional connectivities (SC and FC, respectively). For this purpose, we scanned experienced meditators and control (naive) subjects using magnetic resonance imaging (MRI) to acquire structural and functional data during two conditions, resting-state and meditation (focused attention on breathing). In this way, we aimed to characterize and distinguish both short-term and long-term modifications in the brain’s structure and function. First, we performed a network-based analysis of anatomical connectivity. Then, to analyze the fMRI data, we calculated whole-brain effective connectivity (EC) estimates, relying on a dynamical network model to replicate BOLD signals’ spatio-temporal structure, akin to FC with lagged correlations. We compared the estimated EC, FC, and SC links as features to train classifiers to predict behavioral conditions and group identity. The whole-brain SC analysis revealed strengthened anatomical connectivity across large-scale networks for meditators compared to controls. We found that differences in SC were reflected in the functional domain as well. We demonstrated through a machine-learning approach that EC features were more informative than FC and SC solely. Using EC features we reached high performance for the condition-based classification within each group and moderately high accuracies when comparing the two groups in each condition. Moreover, we showed that the most informative EC links that discriminated between meditators and controls involved the same large-scale networks previously found to have increased anatomical connectivity. Overall, the results of our whole-brain model-based approach revealed a mechanism underlying meditation by providing causal relationships at the structure-function level.

## Introduction

“The mind is definitely something that can be transformed, and meditation is a means to transform it,” wrote in 2015 the world’s foremost Buddhist leader, the 14th Dalai Lama, in his book “The Wheel of Life: Buddhist Perspectives on Cause and Effect”.

Despite having its roots in an ancient Eastern tradition, in the last years, meditation has become increasingly practiced in the Western society, becoming a focus of scientific interest (Ricard et al., 2014; Millière et al., 2018; Hilton et al., 2017; Vieten et al., 2018; Davidson and Dahl, 2018; Afonso et al., 2020).

The term meditation entails all those training practices designed to get aware of mental and bodily processes, which can be clustered into two broader types: concentrative and open awareness practices. The former type requires attention to be voluntarily directed and sustained toward either an internal or external object (e.g., breath-awareness, bodily sensations, musical mantras), whereas the latter implies letting attention opened to whatever comes to the mind.

In line with the above statement from the Dalai Lama, several studies have shown an association between meditation practice and behavioral benefits that result in improvements in attention (Lutz et al., 2008; Valentine and Sweet, 1999), emotional regulation (Miller et al., 1995; Wenzel et al., 2020), and well-being more in general (Peterson and Pbert, 1992; Grossman et al., 2004). In the last decades, different studies have found that experienced meditators show changes in brain morphology compared to matched controls. The first morphometric study conducted by Lazar and colleagues demonstrated that areas involved in interoception and attentional processes, such as the anterior insula and the prefrontal cortex (PFC), were thicker in experienced meditators than controls (Lazar et al., 2005). Since then, several studies investigated meditation-induced brain morphology changes, mainly by measuring cortical thickness (Lazar et al., 2005; Grant et al., 2013; Kang et al., 2013), gray matter volume (Hölzel et al., 2008, 2010; Vestergaard-Poulsen et al., 2009; Pagnoni and Cekic, 2007; Tang et al., 2020), and white-matter integrity (DTI) (Tang et al., 2010, 2015; Luders et al., 2011; Fayed et al., 2013; Posner et al., 2014).

Along with anatomical changes, several cross-sectional studies have found functional changes in experienced meditators compared to controls across large-scale networks, such as the central executive network (CEN), the default mode network (DMN) and the salience network (SN) (Doll et al., 2015; Hasenkamp et al., 2012; Froeliger et al., 2012; Garrison et al., 2015; Kong et al., 2016; Mooneyham et al., 2017; Gard et al., 2014; Irrmischer et al., 2018; Lim et al., 2018). A recent study explored information processing across the whole-brain network, reporting higher dynamical complexity during resting-state in experienced meditators than in healthy controls. At the same time, they found that meditation appears to be characterized as a state of reduced information processing, indicating a switch to a less complex regime compared to the resting state (Escrichs et al., 2019). Similar evidence comes from the work conducted by Toutain et al. (2020), in which they investigated topological stability across relaxation state and meditation in experienced meditators using the electroencephalogram (EEG). The authors demonstrated an increase in stability of global topological patterns during meditation compared to relaxation state (Toutain et al., 2020). Nonetheless, these phenomenological studies cannot by nature provide a mechanistic explanation about how long-term meditation practice shapes the global spatio-temporal BOLD structure via the underlying interregional connections, which instead requires a model-based approach.

Whole-brain computational modeling is one of the most potent tools used to study the link between macroscopic functional dynamics and the underlying structural connectome (Deco and Kringelbach, 2014; Deco et al., 2017; Jobst et al., 2017). They have been mostly used to study the resting-state as well as cognitive functions, including ”consciousness” and alterations thereof. These studies have focused on generating empirical FC with empirical anatomical SC, for psychedelic states (Deco et al., 2018; Herzog et al., 2020; Kringelbach et al., 2020), sleep stages (Jobst et al., 2017; Ipiña et al., 2020) and consciousness disorders (Lopez-Gonzalez et al., 2020; Cofré et al., 2020). In contrast with this work, another line of research has focused on the estimation of the ”effective connectivity”, which describes the directional influence that one region exerts on another in a dynamic model (Friston, 2011). Effective connectivity profiles provide new insights into the causal mechanisms of neuroimaging results by determining the propagation of activity (as a proxy for information) between different areas. Indeed, a challenge of interpreting FC patterns at the whole-brain level is that the BOLD correlation between each pair of areas does not simply follow from connections between them, but also involve network effect via third-party areas. On the biological side, EC estimates represent the modulation of synaptic efficacies due to various factors like neuromodulation, changes in local excitability, etc. that occur in a task-specific manner. A recent approach has developed a model and estimation method, the ‘MOU-EC’, to constrain EC using the SC topology, which forces the model to generate FC by modulating anatomical connections (Gilson et al., 2016, 2020). The MOU-EC model relies on the multivariate Ornstein-Uhlenbeck (MOU) process whose diffusion-type dynamics are used to reproduce the propagation of BOLD signals across the whole brain, and explain them by the structure of the directional EC. Importantly, the EC is masked by the underlying anatomy (DTI tractography), which allow us to evaluate the influence of the measured differences in SC across the two subject groups on the generated BOLD signals. This method has proven capable of extracting biomarkers for cognition as well as subject identification (Senden et al., 2018; Pallares et al., 2018). Moreover, effective connectivity profiles have been proved to be helpful to understand the mechanisms behind brain disease (Adhikari et al., 2020), mental illness (Rolls et al., 2018) and developmental disorders (Rolls et al., 2020).

In this work, we aimed at investigating how meditation-induced changes in the anatomical pathways are associated with the reorganization of the spatio-temporal structure using a dataset consisting of 19 experienced meditators and 19 healthy controls scanned during meditation and resting-state. To do so, we explored whole-brain changes in white-matter tracts following extensive meditation training using a network-based non-parametric approach. We tested whether meditators would show enhanced anatomical connectivity across different large-scale networks. Then, we applied the MOU-EC whole-brain model (Gilson et al., 2016, 2020) in order to relate changes in SC to changes in FC and EC, both across subject groups and across conditions. We compared the effective, functional, and structural connectivity measures as features to train the multinomial linear regression (MLR) and the first-nearest neighbor (1NN) classifiers. Since EC profiles hold information about the underlying structural connectome, we expected EC features to hold more predictive power than FC or SC alone, allowing us to disentangle characteristic information propagation patterns of both groups and conditions.

## Methods

### Participants

A total of 19 experienced meditators and 19 healthy controls were selected from a dataset previously described in Escrichs et al. (2019). In brief, the meditator group was recruited from Vipassana communities of Barcelona, Catalonia, Spain (7 females; mean age=39.8 years (SD=10.29); education=13,6 years; and meditation experience=9.526,9 hours (SD=8.619,8). Meditators had more than 1,000 hours of meditation experience and maintained the daily practice (*>* 1 hour/day). Healthy controls were well-matched participants for age, gender, and educational level, with no previous meditation practice experience (7 females; mean age= 39,75 years (SD=10,13); education=13,8 years). No significant differences in terms of age (p*>*.05), educational level (p*>*.05), and gender (p*>*.05) were found between groups. All participants reported no history of past neurological disorder and gave written informed consent. The study was approved by the Ethics Committee of the Bellvitge University Hospital according to the Helsinki Declaration on ethical research.

### Experimental conditions

Functional data were acquired for two conditions, resting-state and meditation, for a total scanning time of (*≈* 30 min). Participants were asked to stay still in the scanner and move as little as possible throughout the experiment. First, we asked subjects to fixate a cross in the middle of the screen while trying not to think about anything in particular to acquire resting-state data (*≈* 15 min).

Then, we acquired functional data during meditation (focused attention on breathing) (*≈* 15 min). Both groups were asked to practice Anapanasati meditation. Participants had to focus on their natural breathing patterns, trying to recognize whenever their minds were wandering to switch back attention on their breathing. Before the scanning, we instructed control subjects to perform meditation following the instruction given by S.N. Goenka. All subjects confirmed that they understood the procedure before entering the scanner.

### MRI Data Acquisition

MRI images were acquired on a 3T (Siemens TRIO) using 32-channel receiver coil. The high resolution T1-weighted images were acquired with 208 contiguous sagittal slices; repetition time (TR)= 1970ms; echo time (TE)= 2.34ms; inversion time (IT)= 1050ms; flip angle= 9°; field of view (FOV)= 256mm; and isotropic voxel size 1mm. Resting-state and meditation fMRI images were performed by a single shot gradient-echo EPI sequence with a total of 450 volumes (15 min); TR= 2000ms; TE= 29ms; FOV= 240mm; in-plane resolution 3mm; 32 transversal slices with thickness= 4mm; flip angle= 80°. Diffusion-weighted Imaging (DWI) data were acquired using a dual spin-echo DTI sequence with 60 contiguous axial slices; TE= 92ms; FOV = 236mm; isotropic voxel size 2 mm; no gap, and 118×118 matrix sizes. Diffusion was measured with 64 optimal non-collinear diffusion directions by using a single b value= 1,500s/mm2 interleaved with 9 nondiffusion b0 images. A frequency-selective fat saturation pulse was applied to avoid chemical shift misregistration artifacts.

### fMRI preprocessing: resting-state and meditation

Functional MRI images (resting-state and meditation) were preprocessed using version 3.14 of the Multivariate Exploratory Linear Optimized Decomposition into Independent Components (Beckmann and Smith, 2004, MELODIC), which is part of FSL (http://fsl.fmrib.ox.ac.uk/fsl). Images were preprocessed as follows: removal of the first five time-points, motion correction through MCFLIRT (Jenkinson et al., 2002), non-brain removal using the Brain Extraction Tool (Smith, 2002, BET), rigid-body registration, smoothing with 5mm FWHM Gaussian Kernel, and a high-pass filter cutoff set at 100.0s. To discard artifactual components, we applied FIX (Griffanti et al., 2014, FMRIB’s ICA-based Xnoiseifier) using the default parameters to clean the data independently for each subject. Then, the cleaned functional fMRI data were co-registered to the T1 image and the T1 was co-registered to the MNI (Montreal Neurological Institute) space by using FLIRT (Jenkinson and Smith, 2001). The resulting transformations were applied to warp the atlas from MNI space to the cleaned functional data in native space by using a nearest-neighbor interpolation method. Finally, time series in the native EPI space were extracted using fslmaths and fslmeants for 100 cortical using the 7-Networks Schaefer Parcellation (Schaefer et al., 2018) and 16 subcortical regions from the Melbourne subcortical functional parcellation (Tian et al., 2020).

### Probabilistic Tractography analysis

A whole-brain structural connectivity matrix (SC) was computed individually for each subject in their native MRI diffusion space with the same parcellation mentioned above. Analysis was performed using the FMRIB’s Diffusion Toolbox (FDT) in FMRIB’s Software Library www.fmrib.ox.ac.uk/fsl. First, DICOM images were converted to Neuroimaging Informatics Technology Initiative (NIfTI) format using dcm2nii www.nitrc.org/projects/dcm2nii. The b0 image in native space was co-registered to the T1-weighted image using FLIRT (Jenkinson and Smith, 2001), and the co-registered T1 image was co-register to the standard space. The resulting transformation was inverted and applied to warp the atlas in MNI space to the native MRI diffusion space by applying a nearest-neighbor interpolation algorithm. Second, diffusion-weighted images were analyzed using the processing pipeline of the FMRIB’s Diffusion Toolbox (FDT) in FMRIB’s Software Library www.fmrib.ox.ac.uk/fsl. First, non-brain tissues were extracted using the Brain Extraction Tool (Smith, 2002, BET), eddy current distortions and head motion were corrected by using eddy-correct tool (Andersson and Sotiropoulos, 2016), and the gradient matrix was reoriented to correct for subject motion (Leemans and Jones, 2009). Then, crossing fibers were modeled using BEDPOSTX, and the probability of multi-fiber orientations was computed to improve the sensitivity of non-dominant fiber populations (Behrens et al., 2007). Then, Probabilistic Tractography was performed for each subject in native MRI diffusion space using the default settings of PROBTRACKX (Behrens et al., 2007). The connectivity probability SC_ij_ between brain areas *i* and *j* was calculated as the total proportion of sampled fibers in all voxels in brain area *i* that reach any voxel in brain area *j*. Since DTI does not capture fiber directionality, the SC_ij_ matrix was then symmetrized by computing their transpose matrix SC_ij_ and averaging both matrices.

### Network-based statistics

To explore group differences at the anatomical level, we analyzed the whole-brain DTI matrices parcellated into 116 ROIs using the ”Network-Based Statistic Toolbox v1.2 (NBS)” (Zalesky et al., 2010). The NBS, which has been designed to test the hypothesis under the connectome framework, is the network-based equivalent of the suprathreshold cluster-based test (Bullmore et al., 1999; Nichols and Holmes, 2002).

We implemented the NBS to test the hypotheses of structural (SC) over-connectivity (meditators*>*controls) or under-connectivity (meditators*<*controls) following extensive training. A nonparametric permutation approach, with 10000 random permutations, was used to estimate the null distribution of the maximal component size for each of the mentioned hypotheses. At each permutation, values stored in each subject’s SC matrix were used to compute a t-test for each pairwise association contrasting the two groups. Then, we established a primary threshold, *t* = 3 to the t-statistic to determine a set of suprathreshold links from which the connected components and their respective size were identified. The decision to set the primary threshold, *t*, at three for the t-test was based on the fact that we were only interested in a medium or above effect-size, calculated as follows: *t* = *sqrt* (*N*) * 0.5, where *N* is the number of subjects, in our case 38, and 0.5 is the Cohen’s coefficient. Finally, for each contrast (meditators*>*controls and meditators*<*controls), topological clusters of structural links that showed significant differences (p*<*.05) between the two groups were extracted and represented using the Matlab toolbox ”BrainNet Viewer” (Xia et al., 2013).

### Whole-brain MOU-EC model and parameter estimation

As described in Figure 1, the dynamic generative model MOU-EC was used to obtain whole-brain connectivity estimates from the datasets parcellated into 116 ROIs as previously described. First, we used the SC matrices extracted through the probabilistic tractography analysis (black and white matrix at the left top of Fig.1) to constrain the model’s topology by setting a threshold to retain a 30% density of anatomical pathways. For within-group comparisons, we extracted two probabilistic SC matrices, one for meditators and one for controls, by computing the average SC of participants of each group. In contrast, to compare groups within each task, we converged the overlapping links that were above threshold in both groups, by intersecting the SC matrix of meditators with the SC matrix of controls. For each fMRI session, the BOLD autocovariance was calculated, both with and without time lag (blue FC0 and green FC1 matrices in Fig. 1), and then reproduced by the model. Before calculating autocovariance matrices, fMRI data were further filtered with narrowband 0.04-0.07 Hz to avoid artifacts (Glerean et al., 2012). This framework relies on a dynamic system with linear feedback to extract spatio-temporal information about the BOLD dynamics and directed connectivity estimates (i.e, EC), namely the Multivariate Ornstein-Uhlenbeck (MOU) process. The MOU process, analogous in continuous-time of the discrete-time multivariate autoregressive process, can be mathematically described as follows:

**Figure 1:**
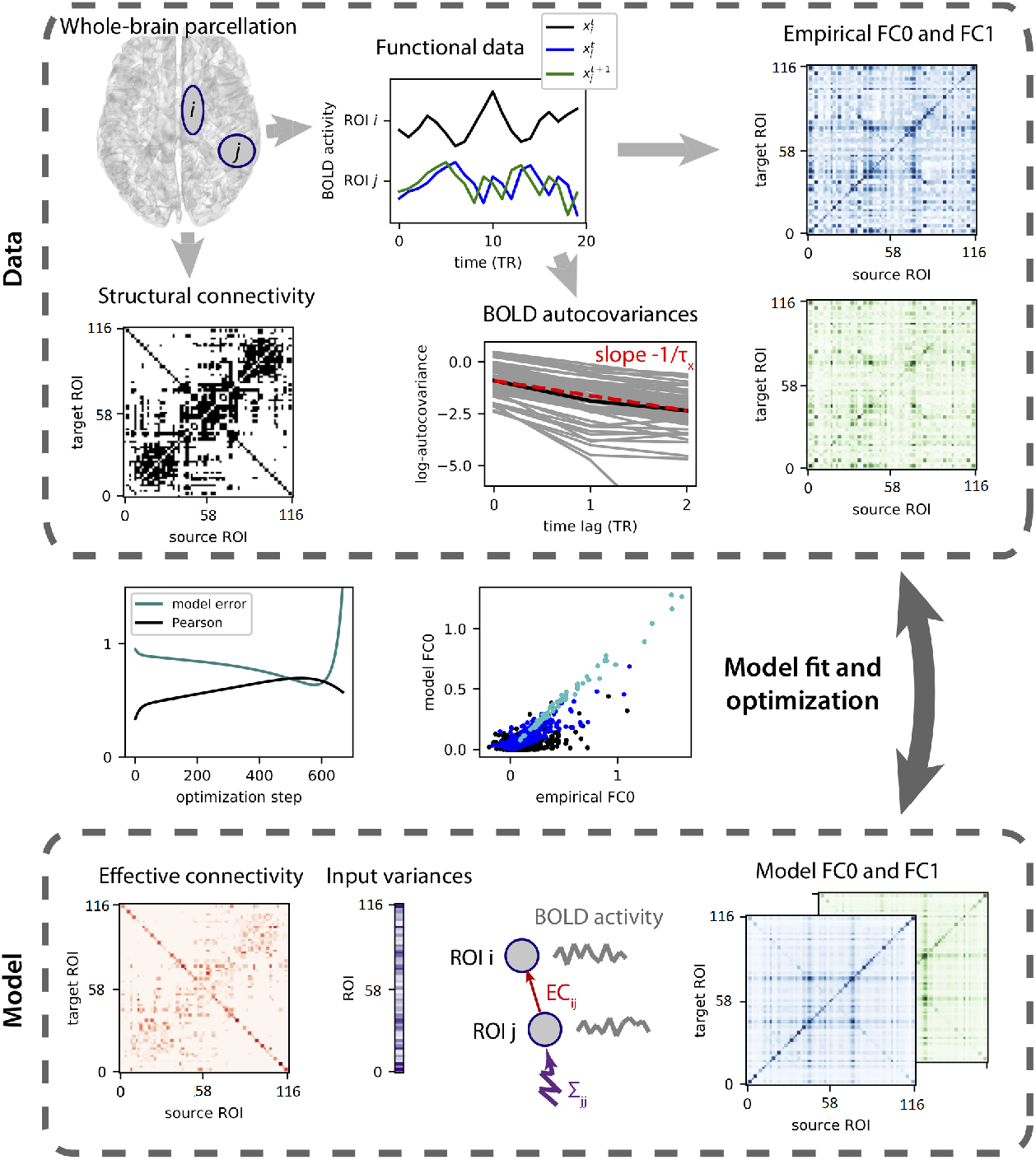
Model parameters estimation (adapted from Gilson et al., 2020). To capture BOLD dynamics, the model uses the time-series parcellated into 116 ROIs (box on the left top) to calculate two autocovariance matrices (FC0 and FC1 matrices) both with and without time lag (blue and green lines in the central box on top). The SC connectivity matrix (black and white matrix on the left) is used as a binary matrix to constrain the model’s topology to existing connections. Besides the effective connectivity estimates (Pink matrix at the left bottom), the autocovariance matrices (green and blue matrices at the right bottom) are also reproduced by the dynamic model. The model undergoes an optimization procedure so that, at each step, the estimated FC matrices are evaluated with regards to the empirical FC0 and FC1 matrices. The optimization steps are repeated until reaching a high Pearson correlation coefficient between the model’s and the empirical FC matrices, reducing this way the model error (depicted as the green and black lines in the central box on the left of the figure).

**Figure 2:**
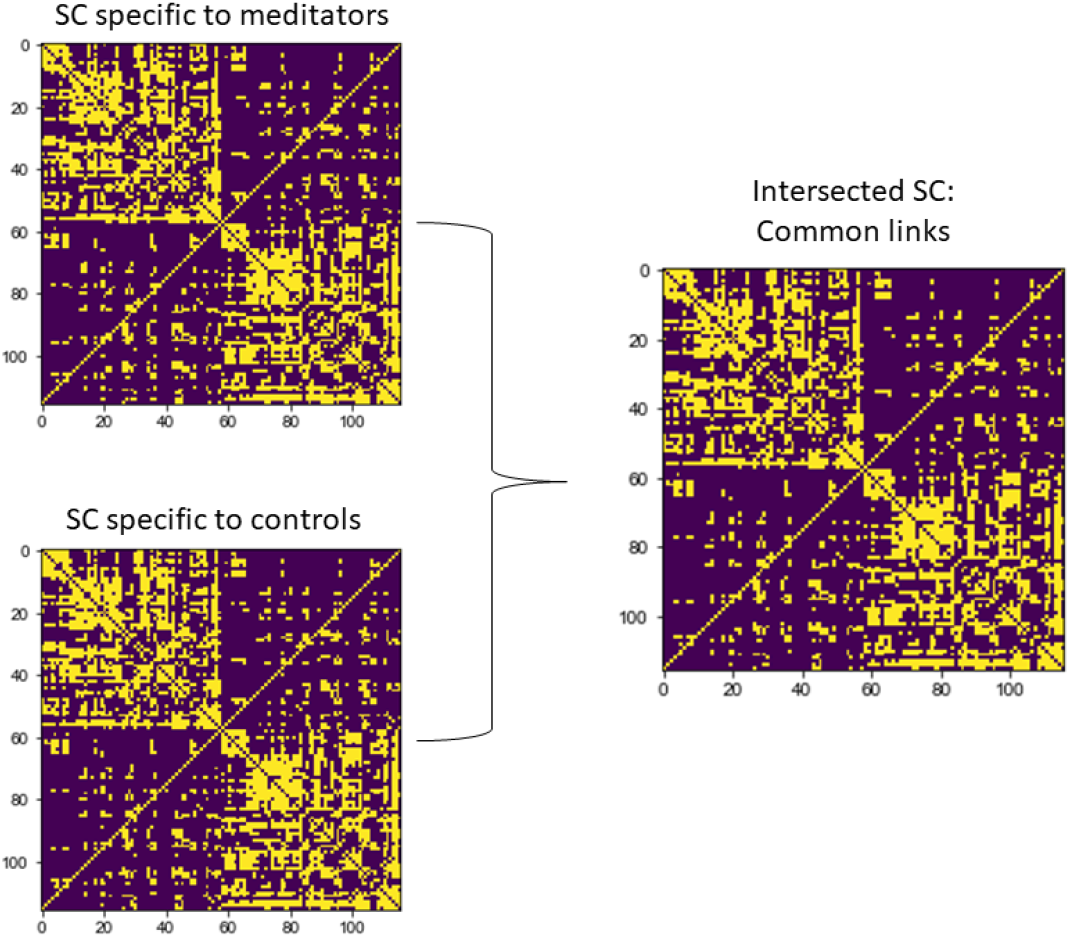
Intersected SC matrix for each group. To estimate functional dynamics of both groups during each condition, we intersected the SC matrix specific to each group (two matrices on the left side) and we generated a new SC matrix with all the links above-threshold common to both meditators and controls (SC matrix on the right side).

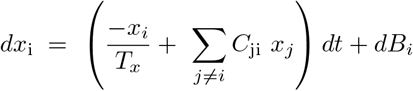

Where *x*_i_ denotes the activity of ROI *i*, which is influenced by the activity of other nodes and decays exponentially by the time constant *T*_*x*_. The information about the direct connection between ROIs(*i,j*) (i.e., EC) is stored in the matrix *C*_ji_ (pink matrix at the left bottom of Fig.1), whose skeleton is determined by the SC matrix so that weights for nonexistent connections are kept to 0 while those for existing links are estimated from the FC matrices. The variable *dB*_*i*_ refers to independent fluctuating inputs (i.e., local variability), consisting of the diagonal covariance matrix Σ (vector of variances at the bottom of Fig.1). The estimated FC is determined by the propagation of the local variability generating network feedback via the EC. The model’s parameters Σ, and *C*_ji_ undergo an iterative gradient-descent optimization procedure such that the model is tuned to reproduce the empirical FC0 and FC1 with the minimal error and the maximum values of Pearson correlation coefficient mean for each session. For further mathematical details, we refer the readers to Gilson et al. (2016, 2020). The code related to model optimization and classification is based on the open-source language python, and it is available at github.com/MatthieuGilson/WBLEC_toolbox.

### Classification based in EC/FC weights

The classification procedure was run using the scikit-learn package (Pedregosa et al., 2011) based on Python language. We performed four binary classification types: classification of conditions (resting-state vs. meditation) within each group (controls and meditators, separately) and then group classification (control group versus meditators), one for each state. To construct the feature arrays, we used the probabilistic SC matrix together with model’s FC and estimated EC matrices. First, we vectorized the SC, FC and EC matrices for each session (e.g., resting-state or meditation). To reduce dimensionality, we selected the lower triangle of the symmetric SC and FC matrices, resulting in a vector of 6670 SC links and one of 6670 FC links for each session. We applied the SC mask specific for each group, or the intersected matrix between the two groups, to the EC matrix in order to extract the EC vectors. Then, we z-scored within each session the SC, FC and EC links using the mean and standard deviation of the corresponding vectorized connectivity measures. Therefore, for each session of each subject, here referred to as sample, we had the ranking of the vectorized elements of EC, FC, and SC as features to train two different classifiers, namely the 1-Nearest Neighbor (1NN) and the Multinomial linear regression (MLR). This choice is because the two algorithms capture the data’s different properties. Indeed, for the 1NN, we used Pearson correlation as a metric to evaluate the inverse distance between samples (i.e., vectorized connectivity matrices). In this way, the 1NN classifier predicts the test-set session’s class by identifying the trainset’s most similar session. On the other hand, the MLR is a supervised learning algorithm for high-dimensional data optimal for linear classification. In order to predict the class (i.e., group or condition), the model regressors are adjusted. The MLR tunes weights for each dimension of the input features, allowing efficient feature selection procedures.

To train and cross-validate the two classifiers (1NN and MLR), we used 80% of the datasets for training the algorithms and the remaining 20% for testing. We repeated the random splitting procedure 50 times to assess the impact on the performance of different splits of data in train/test sets. Only the accuracy of the 50 predictions on unknown data from the test-set were considered to evaluate models’ performance.

We compared accuracy distributions using the Wilcoxon rank-sum method understand which combination of classifier and metric performed better. Moreover, to investigate whether classification results were significantly above chance, we compared accuracy distributions of real-labeled data with surrogate data using the Wilcoxon rank-sum test.

### Signatures extraction: support networks

We applied the Recursive-Feature Elimination with the MLR to extract group-specific signatures. This algorithm is largely used in machine learning to rank the features according to their relevance for the classification (Guyon et al., 2002). RFE applied to MLR allows to iteratively select a subset of features by pruning at each iteration the least important features from the whole set until only the most relevant links are left. We applied the RFE algorithm to group classification during meditation and resting-state, separately, to disentangle the EC links (akin, support network) specific to the group. For each application, we randomly selected 80% of data to train and fit the MLR using RFE. Simultaneously, the remaining samples were used to evaluate the accuracy of MLR using a different number of features based on the order given by the RFE ranking. We repeated this cross-validation procedure ten times, randomly selecting 80% of samples for the train set and 20% for the test set. Finally, we chose a subset of features for which the classifier performance was stable across iterations and including additional features did not provide a significant increase in classification accuracy.

## Results

### White-matter changes induced by long-term meditation practice: NBS results

We used the NBS toolbox for identifying network connectivity differences at the anatomical level following extensive training. We compared probabilistic DTI matrices of experienced meditators and controls, and tested for both increased (meditators*>*controls) and decreased (controls*>*meditators) probability of white-matter connectivity.

Significant results of nonparametric statistical analysis on anatomical matrices using the NBS toolbox are shown in Figure 3. When testing the hypothesis of increased WM connectivity following extensive training (meditators*>*controls), we found a component of 9 edges and 10 nodes, mainly involving interhemispherical connections, that was significantly enhanced in meditators compared to controls (p= .009).

**Figure 3:**
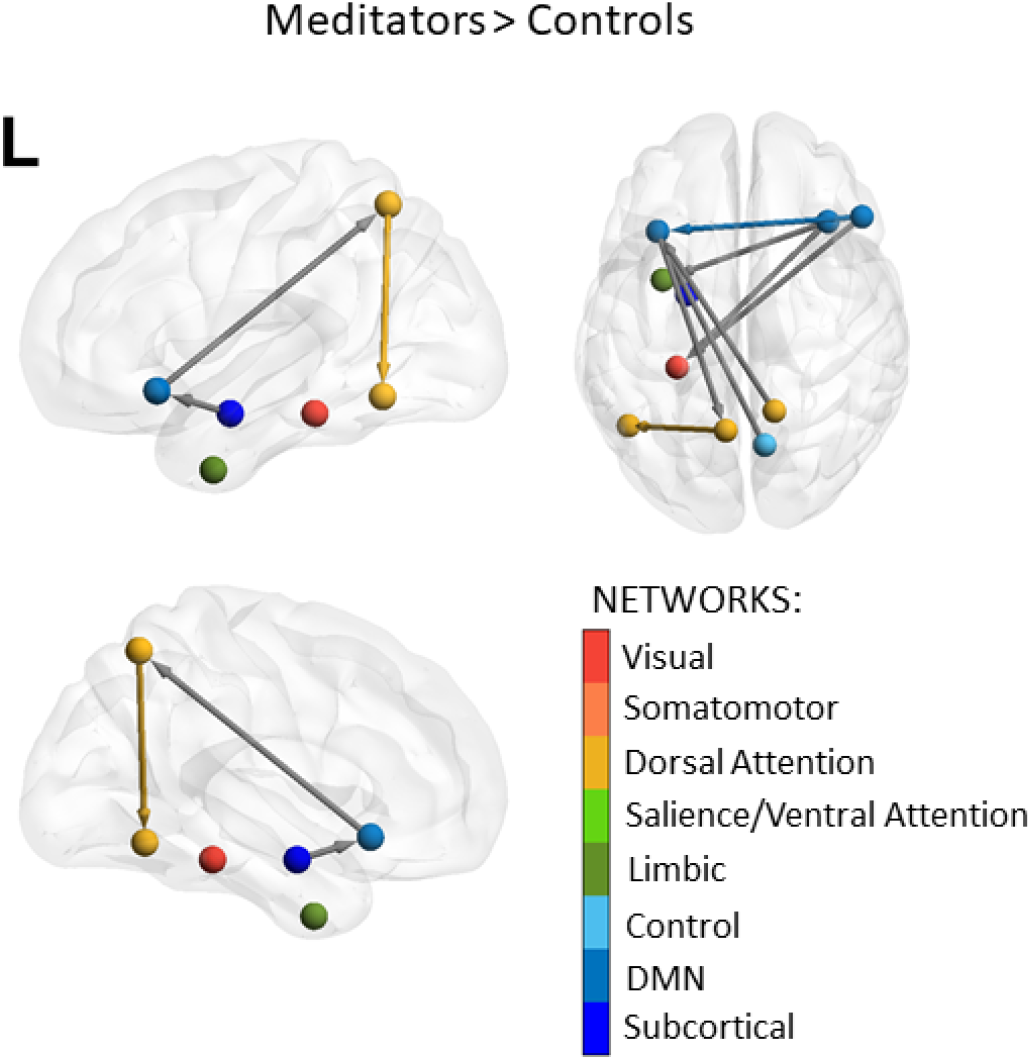
Increased network-related SC in the meditators group compared to the control group. The significant network resulting from the contrast meditators*>*controls, comprising 9 edges and 10 nodes, is represented from a lateral (left top brain), medial (left bottom brain), and dorsal (right bottom brain) views. Nodes are colored according to the hue of the network they belong. Connections within the same network nodes are presented with the color hue of that specific network, while a gray arrow represents edges between nodes of different networks. The two yellow nodes correspond to the left posterior area 1 and area 5 of the dorsal attention network, the green node refers to region 1 of the limbic temporal pole, and the red node represents the visual area 1. The amygdala, part of the subcortical network, is represented by the dark blue node, while the three lighter blue nodes refer to the prefrontal cortex (PFC) areas belonging to the default mode network.

The component comprised regions belonging to the dorsal attention, the default-mode (DMN), the control, and the limbic networks. In particular, we found an increased probability of connection in the meditator group within and between the posterior dorsal attention network and prefrontal regions belonging to the DMN. The left prefrontal cortex (PFC) area of the DMN was also found to show increased connectivity with three areas in the contralateral hemisphere: the precuneus, part of the control network, the portion of ventral PFC belonging to the DMN, and the right amygdala. Furthermore, in the meditator group, we found that two DMN nodes in the right PFC had increased connectivity with the left visual area and the left limbic temporal pole. On the other side, for the contrast controls*>*meditators, we obtained no significant results in component extent (p = .72), suggesting that meditators do not show any network-related decrease in anatomical connectivity compared to controls.

### Classification of conditions within each group

Next we investigated functional changes, as quantified by FC and EC, with the aim to relate them to the SC changes. For EC, we fitted the dynamic whole-brain model informed by the SC for the two groups of subjects, meditators and controls. We investigated using machine-learning algorithms whether meditation condition would be accurately differentiated from resting-state in meditators and in controls subjects who were naive to meditation practice. Average accuracies corresponding to the classification of conditions (i.e., meditation and resting-state) within each group following the described feature extraction and validation methods are shown in Figure 4. Moreover, we compare the distributions of accuracies for different features or classifiers using the Wilcoxon rank-sum test, reporting p-values.

**Figure 4:**
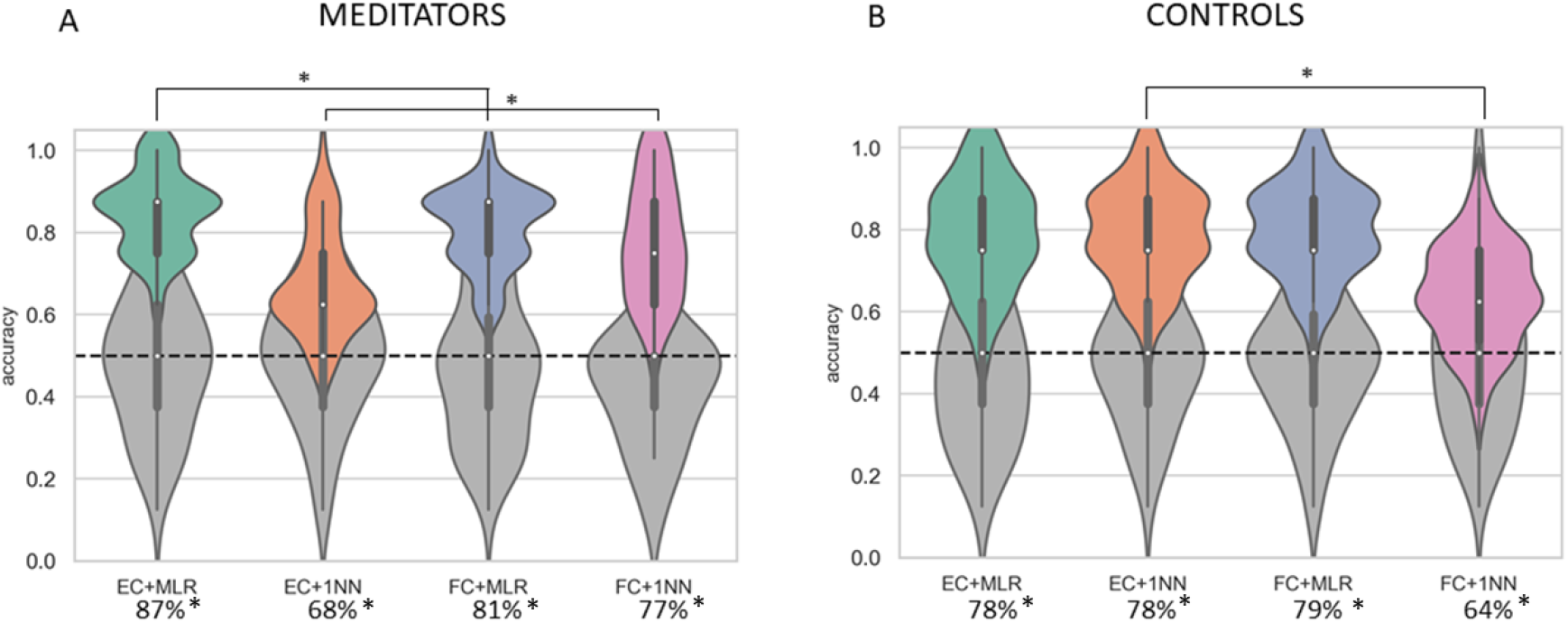
Resting-state vs. meditation: classification accuracies within each group. Accuracy distributions are shown for meditators (left panel) and controls (right panel) The violin plots in green show the accuracies for the MLR classifier using the features related to EC, while the blue ones represent MLR performance using FC features. Accuracies related to 1NN are shown in orange for EC and in pink for FC features. For both groups, all the combinations between classifiers (MLR and 1NN) and metrics were significantly higher than chance-level (p*<*.0001).

For features extracted with the estimated EC, we reported a significantly higher performance with an average increase in accuracy of 6% compared to FC features for the MLR classifier in the meditator group (Fig. 4A) (p=.004). Differences in MLR performance, depending on the type of features, were not significant (p*>*.05) for the control group (Fig. 4B). Performance of 1NN for distinguishing resting-state from meditation in the meditator group appeared to improve when using FC features compared to EC (p*<*.0001), indicating that FC patterns are overall more similar between the two conditions, while differences in EC measures are more localized. However, MLR significantly outperformed the 1NN, both when using EC (p*<*.0001) and FC features (p*<*.0001) for the meditator group. This suggests that the it is less the global EC (captured by the Pearson correlation as a similarity measure for the 1NN) profile than specific EC links that significantly vary across conditions in the meditator group. In contrast, EC significantly improved 1NN accuracy for the control group, with a performance drop of 14% on average using FC (p*<*.0001) while the MLR performance was not affected by the type of features for control subjects (p*>*.05). Nevertheless, the performance of classifiers was highly above chance-level both for meditators and controls (p*<*.0001) using all metrics and classifiers.

These results showed that meditation induces characteristic connectivity patterns in brain activity which can be differentiated with high precision even in control subjects naive to the meditation practice.

### Group classification within each condition and signatures extraction

To further investigate the relationship between the impact of extensive training on white-matter connectivity with the whole-brain functional dynamics, we classified the groups (meditators vs. controls) separately in the two conditions (meditating and resting). Following the differences in SC across the two groups in Fig 3, we also included a third set of features consisting of the vectorized SC matrix specific to each subject. The group classification results for each condition are shown in Figure 5.

**Figure 5:**
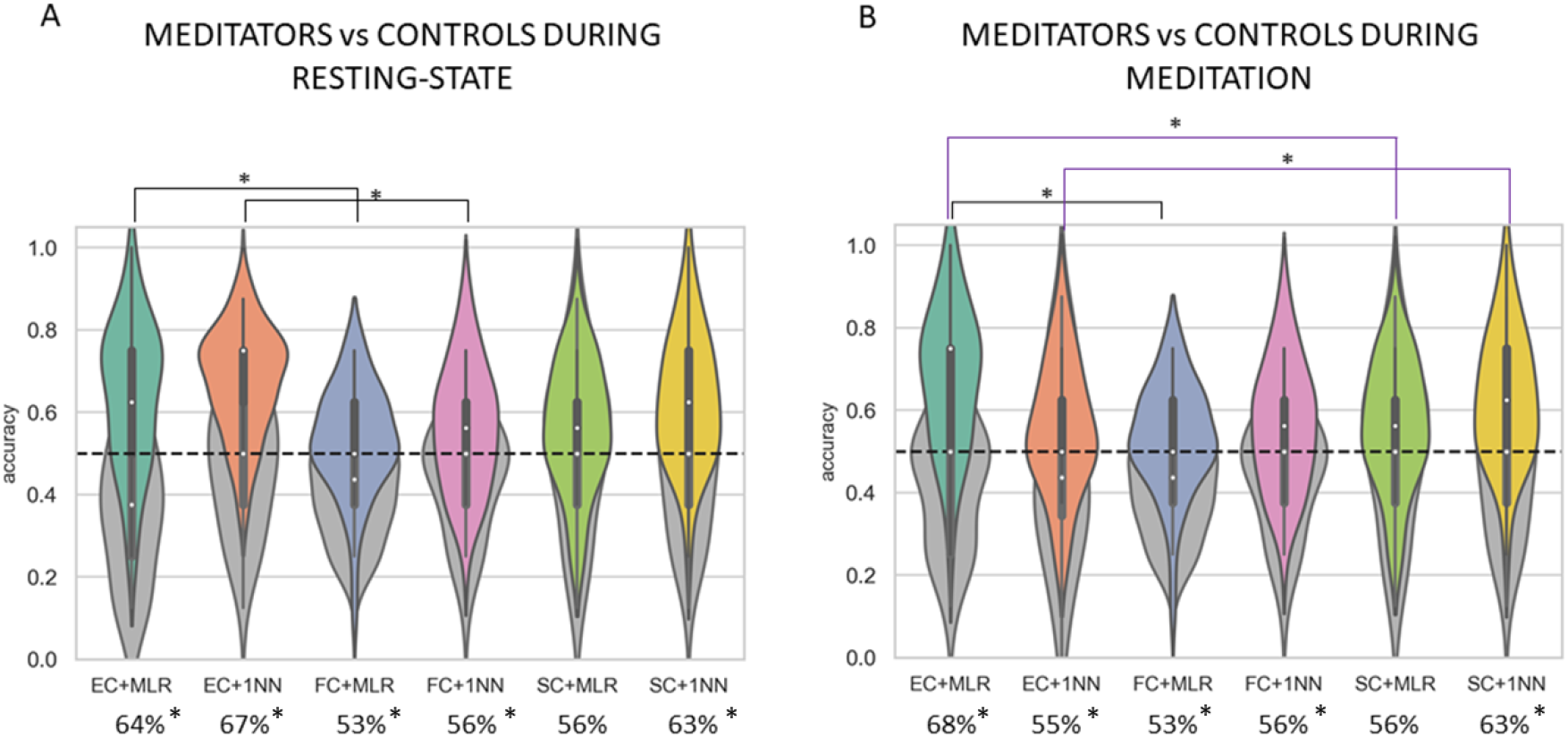
Results of group classification during meditation and resting-state. The performance of 1NN and MLR in discriminating the two groups are shown for resting-state condition (A) and meditation task (B). Significant differences in performance depending on the type of features are shown in black for the contrast EC-FC features and in purple for the contrast EC-SC features. Violin plots colored in gray represent accuracy distributions of surrogate data. EC features led to the highest average accuracies with either the MLR or the 1NN, depending on the condition. While the 1NN yielded the best performance for the resting-state sessions (average accuracy of 67%), the MLR outperformed the 1NN when comparing the two groups during the meditation task (average accuracy of 68% against 55%, respectively).

The results of classification using resting-state sessions (Fig.5A) suggested that the better combination of classifier and metric is the 1NN with EC features, with an average accuracy of 67%. The 1NN appeared to be better suited for discriminating meditators and controls in rest, suggesting that data are overall clustered and that samples with the same labels are closer (i.e., showing similar EC profiles). In particular, we found significant higher accuracy using EC features compared to FC features in the group classification during resting-state, both for the MLR (p=.01) and the 1NN (p=.002). EC features also yielded higher accuracies for the MLR compared to FC features, but the difference did not reach significance (p=.06). There were no difference in performance between FC and SC, both when using the 1NN (p*>*.05) and the (p*>*.05), nor between EC and SC when using the 1NN (p*>*.05).

When comparing the two groups while meditating (Fig. 5B), we found that the MLR showed better performance (average accuracy= 68%) than the 1NN (average accuracy=55%) when using EC features. This indicates that differences between the two groups while meditating are based on specific weights of some EC features rather than the overall EC profile. Again, EC features yielded to a better MLR performance compared to FC (p*<*.0001), and compared to SC (p=.005). In contrast, we found a significant increase in 1NN performance when using SC features compared to EC (p =.03). Difference in 1NN performance when contrasting FC and SC features were not significant (p*>*.05).

These results suggested that meditation condition is in general better to distinguish between subjects of the two different groups, as compared to resting-state. Moreover, EC features appeared to be more informative than FC and SC in discriminating the two groups, both when resting (using MLR and 1NN) and meditating (using MLR). This indicates that the combination of information of SC and FC by the anatomo-functional whole-brain model better characterizes the differences between the two groups. Note that the performance of combination of classifiers (1NN and MLR) and metrics (EC,FC,SC) was significantly higher than change-level (p*<*.05) for all cases except the combination MLR with SC features which did not reach significance (p*>*.05).

Following the results of classification performance, we performed Recursive-Feature Elimination (RFE) with the MLR classifier during meditation and resting-state conditions, separately. We investigated which EC links contributed the most to the accurate distinction between subjects of the two groups. Results of RFE for group comparison during meditation are presented in figure 6. We found that features relevant for the group classification during meditation were less (15 edges) than those of the resting-state condition (23 edges). In both cases, the support networks showed a prominence of intermispheric connections and connections within the left hemisphere although they were distributed and across frontal and posterior regions. The most relevant features extracted during meditation mostly involved top-down regulation between high-level functioning areas belonging to the control, the dorsal attention, the salience/ventral attention, the DMN and the somatomotor networks. In contrast, the support network of the resting-state condition showed a larger involvement of the visual network and subcortical structures (including the thalamus, the amygdala and the putamen), together with large-scale networks. In particular, the precuneus/posterior cingulate cortex areas belonging to the DMN were found to exert direct influence on nodes belonging to the visual, salience/ventral attention and the posterior dorsal attention network. Moreover, the left PFC areas belonging to the DMN showed a top-down regulation over the putamen together with projections to somatomotor and control networks. At the same time, the left PFC section of the DMN received direct influence from the frontal operculum/insular region belonging to the salience/ventral attention network. This areas was also found to receive top-down projections from the orbito-frontal cortex, part of the limbic network. Connections going from and to the frontal operculum/insular region were also found to be relevant when using meditation sessions. In fact, this area appeared to be modulated by the orbito-frontal cortex, together with the left PFC part of the DMN and the lateral PFC part of the control network.

**Figure 6:**
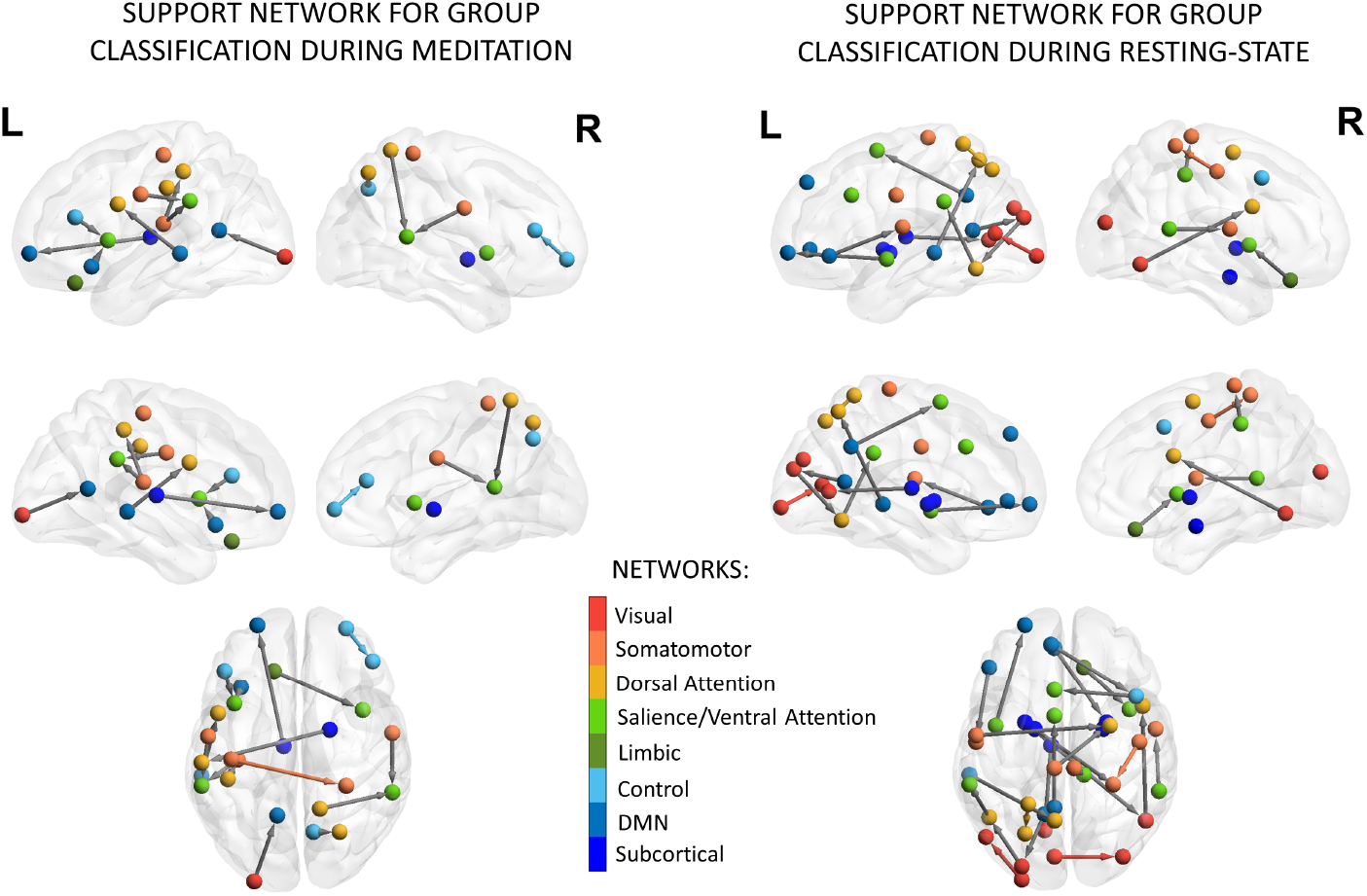
RFE results: Support network for group classification during meditation. The group signatures (i.e., highest-ranked EC features) are presented for the two conditions in sagittal, medial, and dorsal views, both for the right and left hemispheres. The color of the nodes is determined by their belongingness to one of the eight resting-state networks (red= visual; orange= somatomotor; light orange= dorsal attention; light green = Salience/Ventral Attention; dark green = limbic; light blue= control; blue = DMN; dark blue = subcortical). The networks are drawn from the parcellation in 100 ROIs comprising 7 functional divisions (Schaefer et al., 2018) together with 16 subcortical regions (**?**). Gray arrows show the directionality of connection (EC-links) between ROIs of distinct networks. In contrast, direct links between nodes belonging to the same network are colored according to network-specific hue. Visualization of results was generated using the BrainNet Viewer Toolbox (Xia et al., 2013)

Together these results suggest that there are distributed differences between meditators and control subjects in information propagation across large-scale networks, which are more prominent in the left hemisphere.

## Discussion

The purpose of this work was to contribute to research in contemplative neuroscience by shedding light on the causal mechanisms underlying long-term meditation practice. First, we computed a network-based analysis of whole-brain structural connectivity to understand how extensive training modulated anatomical pathways. We showed that meditators had increased white-matter connectivity across several large-scale networks compared to controls. Then, we applied computational modeling in order to understand how differences in the underlying anatomical connectivity were reflected in the brain’s information propagation. We estimated through a model-based approach the whole-brain EC profiles of meditators and controls, using their underlying anatomical connectivity to explain functional dynamics during rest and meditation. Using machine-learning tools, we demonstrated that EC features led to higher performance than FC or SC alone when classifying groups and conditions, indicating that there is a synergy between structural and functional connectivity patterns that can be captured through EC measures. We demonstrated, by applying a feature selection procedure, that the two groups showed differences in information propagation across the same large-scale networks found to have increased SC, both during meditation and resting-state.

The consistency of these findings supports the structure-function mutual relationship hypothesis, according to which there is a high topological correspondence between structural and functional connectivity (Straathof et al., 2019). In fact, we showed that extensive meditation training leads to a structural reorganization of white-matter pathways. We opted for a network-based approach since we were more interested in investigating the effects of long-term meditation practice on interconnected subnetworks rather than focal effects. Using the network-based statistics (NBS) method, we found that experienced meditators, compared to the control group, showed enhanced SC in a subnetwork mostly involving distributed inter-hemispheric connections between areas of the DMN, the dorsal attention, the limbic, the visual, and the control networks.

The second reason is simply a matter of power—the NBS can offer substantially greater power in the right circumstances, which is advantageous in the context of the graph model due to the massive number of multiple comparisons that arise when the hypothesis of interest is tested at every connection. The component highlighted by NBS analysis demonstrated strengthened connectivity between areas involved in attentional, self-referential, and emotional processes. In particular, changes in anatomical connectivity following extensive practice were more prominent between hemispheres and within the left hemisphere (Fig.3), with increased connectivity between the posterior areas of the dorsal attention and the left PFC, part of the DMN. The left PFC area belonging to the DMN appeared to be a central hub, showing increased connectivity also with posterior areas of the control network, and subcortical structures, such as the right putamen and the right amygdala. Further interhemispheric WM tracts that were found to be enhanced in meditators compared to controls regard connections between the left visual area and the right ventral PFC, part of the DMN, which was also observed to have increased connectivity with the limbic temporal pole and the left DMN portion of the PFC. The structural neuroplasticity that we found involved areas whose functions is associated with largely reported meditation-induced behavioral effects, such as improved emotional regulation and reduced stress (Chiesa and Serretti, 2009; Chung et al., 2012; Tang et al., 2016), enhanced attentional skills (Valentine and Sweet, 1999; Brefczynski-Lewis et al., 2007; Semple, 2010; MacLean et al., 2010), and reduced mind-wandering (Brewer et al., 2011).

In line with these results, we found that direct connections (i.e., EC links) between areas of the same large-scale networks that showed increased SC were also part of the support networks that allowed discriminating meditators from controls during resting-state and meditation. In fact, we demonstrated through group classification that meditators could be accurately distinguished from control subjects both during rest (MLR average accuracy =64%) and meditation (MLR average accuracy =67%), and that the maximum accuracy was reached when using EC features. In both meditation and resting-state, the group signatures extracted through RFE were distributed across the cortex but showed a prominence of the left hemisphere and involved EC links between nodes belonging to the DMN, the dorsal attention, the salience/ventral attention, the control, the visual and the somatomotor networks. Notably, the support network using resting-state sessions was langer and showed a bigger involvement of visual and subcortical areas, such as the amygdala, the thalamus and the putamen, compared to the one of meditation condition. Additionally, the frontal DMN nodes appeared to exert direct influence over the control, salience/ventral attention and dorsal attention regions during resting-state. Moreover, we found an upregulation from the frontal operculum/insular region belonging to the salience/ventral attention network sending projections to the PFC area belonging to the DMN during resting-state. The frontal operculum/insular region appeared to play a central role in differentiating between meditator and control groups during resting-state as well as meditation. In both cases, we found that the frontal operculum/insular region received projections from the lateral PFC (part of the control network) and the orbito-frontal cortex (part of the limbic network). This is consistent with the hypothesis that the insular regions play a central role in switching between different networks (Sridharan et al., 2008) and for interoceptive and emotional awareness (Simmons et al., 2013), which is a central skill trained through meditation (Lutz et al., 2008). Although here we went beyond FC by capturing the direct influence that one region may exert on others through EC, these results are in line with previous literature on meditation-induced functional connectivity changes that have found increased coupling for meditation practitioners compared to matched control subjects within and between nodes of dorsal attention network and areas of DMN, and salience networks (Froeliger et al., 2012), as well as connections among the nodes of dorsal attention network, executive, and visual circuits (Kemmer et al., 2015; Boccia et al., 2015).

Despite the differences between the two groups in EC links highlighted by the support network of group classification, we found that it was possible to discriminate resting-state from meditation with high precision also for control subjects naive to meditation practice. The results of condition classification within each group showed that differences between resting-state and meditation for control subjects could be captured with high accuracy both when using EC (MLR average accuracy =78%, 1NN average accuracy =78%) and FC features (MLR average accuracy =79%, 1NN average accuracy =64%). The significant difference in 1NN performance between EC and FC features suggest that the overall EC profile of control subjects is more informative than their global FC profile in discriminating the two conditions. In contrast, in the meditator group the best performance was reached when using the MLR classifier with EC features (average accuracy =87%), indicating that for meditators the differences between the two conditions rely on specific EC links rather than on the overall EC profile.

The present study could be improved in the future in several manners. First, the NBS method does not allow for interpretation at the single edge level since it provides only information about the network behavior as a whole (Zalesky et al., 2010). Therefore, we could not directly relate at the node level the results of SC with the results of RFE highlighting the most informative EC links. An analysis at the link level requires more statistical power, hence many more subjects than those analyzed here. Another limitation of our cross-sectional study lies in the statistical relationship between the structural and functional changes, which could be studied in more depth via a longitudinal experimental design to jointly measure the structural and functional changes induced by meditation practice over a long training period.

## Conclusions

The present work focused on unraveling the anatomical mechanisms and their relationship to brain function underlying long-term meditation practice. We demonstrated the advantage of using a model-based approach to extract effective connectivity estimates compared to standard correlation analysis of BOLD signals, to highlight differences in information propagation between experienced meditators and controls. Moreover, network-based analysis of anatomical pathways revealed a cluster of edges, comprising areas involved in attentional, emotional, self-referential and control processes, for which structural was enhanced in experienced meditators. Effective connectivity links between nodes of the same large-scale networks were also found to be the most discriminative features for distinguishing meditators and control subjects in resting-state and during meditation. Together these results suggest the presence of meditation-induced neuroplasticity at the function-structure level.

## Notes

### Competing Interest Statement

The authors have declared no competing interest.

